# Physical confinement impacts cellular phenotype within living materials

**DOI:** 10.1101/2020.03.25.004887

**Authors:** Hans Priks, Tobias Butelmann, Aleksandr Illarionov, Trevor G. Johnston, Christopher Fellin, Tarmo Tamm, Alshakim Nelson, Rahul Kumar, Petri-Jaan Lahtvee

## Abstract

Additive manufacturing allows three-dimensional printing of polymeric materials together with cells, creating living materials for applications in biomedical research and biotechnology. However, understanding the cellular phenotype within living materials is lacking and a key limitation for their wider application. Herein, we present an approach to characterize the cellular phenotype within living materials. We immobilized the budding yeast *Saccharomyces cerevisiae* in three different photocross-linkable triblock polymeric hydrogels containing F127-bis-urethane methacrylate, F127-dimethacrylate, or poly(alkyl glycidyl ether)-dimethacrylate. Using optical and scanning electron microscopy, we showed that hydrogels based on these polymers were stable under physiological conditions, but yeast colonies showed differences in the interaction within the living materials. We found that the physical confinement, imparted by compositional and structural properties of the hydrogels, impacted the cellular phenotype by reducing the size of cells in living materials compared with suspension cells. These properties also contributed to the differences in immobilization patterns, growth of colonies, and colony coatings. We observed that a composition-dependent degradation of polymers was likely possible by cells residing in the living materials. In conclusion, our investigation highlights the need for a holistic understanding of the cellular response within hydrogels to facilitate the synthesis of application-specific polymers and the design of advanced living materials in the future.

## Introduction

Three-dimensional (3D) printing of natural and synthetic materials for biomedical and biotechnology applications is a promising research field with applications that include screening tools and production platforms in a sustainable economy.^1^ Self-assembling block copolymer hydrogels have been demonstrated for extrusion-based 3D printing, and offer exciting opportunities to create synthetic polymer hydrogel networks that can immobilize microbial cells and recapitulate the environment of a biofilm.^2,3^ These microbe-laden hydrogels form living materials (LMs) that are permissive for metabolic activity and can provide significant improvement with respect to robustness, reproducibility, and scale-up over traditional immobilization methods using natural biopolymers.^4^ The multiscale properties of hydrogels of such polymers allow their applications in diverse fields, such as drug delivery^5^, tissue engineering^6^, and biotechnology^4,7^. Precise material deposition, together with a high degree of spatial control, allows the manufacturing of pre-designed and custom-made structures.^8,9^ One prominent triblock copolymer hydrogel for extrusion-based printing is based on Pluronic F127, which embodies dual-responsive properties towards temperature (sol at 4 °C, gel at 25 °C) and the applied shear forces.^10^ This ABA triblock copolymer, wherein the ‘A’ blocks are hydrophilic poly(ethylene oxide) and the ‘B’ block is a hydrophobic poly(propylene oxide), can self-assemble to form micelles in aqueous solution. As the concentration of the F127 in solution increases, the polymer reaches a critical gel concentration. The Nelson group recently developed BAB triblock copolymer hydrogels for direct-write extrusion printing with hydrophobic poly(alkyl glycidyl ether) ‘B’ blocks that flank a central poly(ethylene oxide) ‘A’ block that exhibits similar stimuli-responsive behaviors as F127.^11^ In contrast to F127, the BAB triblock copolymers form reverse flower micelles in solution.^11–13^ Furthermore, the chain-end modification of BAB and ABA triblock copolymers allows for cross-linking by means of photo-initiated polymerization while or after completion of the 3D printing process to afford robust hydrogel structures.^14–16^ Polymer hydrogels based on F127-dimethacrylate (F127-DMA), F127-bisurethane methacrylate (F127-BUM), and poly(isopropyl glycidyl ether-*stat*-ethyl glycidyl ether)-*block*-poly(ethylene oxide)-*block*-poly(isopropyl glycidyl ether-*stat*-ethyl glycidyl ether) dimethacrylate (PGE-DMA) have previously been reported for encapsulation and direct-write extrusion printing of microbes.^4,17–19^ In all of these cases, the hydrogels maintained the viability and metabolic activity of yeast or bacteria to afford immobilized bioreactors with longterm metabolic activity.^4,17,18^

Methods for the characterization of the physico-chemical properties of such hydrogels, particularly their stiffness, swelling ratio, and rheology, are well established.^20,21^ However, similar robust analysis methodologies for understanding cellular phenotypes of microbial cells confined within hydrogels are lacking, but necessary, before LM-based technologies could be used in specific, reproducible, and efficient processes. Previously, optical microscopy (OM) and scanning electron microscopy (SEM) have been used to investigate cell-gel-morphology^22–24^ and hydrogels themselves^24^–^26^, but only to an illustrative extent. For this reason, we focused on these reliable and accessible microscopy tools and techniques for the characterization of LMs and for the investigation of cellular phenotypes in a physiological environment. In all instances, we used the budding yeast *Saccharomyces cerevisiae*, which has been previously reported to be viable in these materials and assigned the generally recognized as safe (GRAS)^27^ status making it applicable in food and pharma industries.^4,17,18^ In our study, we selected three different functionalized triblock copolymers: F127-DMA^4^, F127-BUM^17,19^, and PGE-DMA^18^. These polymers are advantageous over calcium alginate for microbial encapsulation because the materials are covalently cross-linked and charge-neutral. The carboxylate groups of alginate have previously been shown to inhibit the transport of charged molecules through these hydrogel matrices.^17^ We investigated the stability and degradation of the hydrogels of these polymers after cultivation in a physiological environment, the polymer-cell interface, the localization of cells, the proliferation of colonies, the effect of cellular growth on the polymers, and the effect of physical confinement on cellular phenotype using both OM and SEM methods. Further, we used a computational approach for SEM image analysis to determine cell size changes in living materials and suspension cell cultures. This allowed us to assess the effects of different polymers on the cellular phenotype. The detailed workflow of our study is illustrated in Figure 1.

**Figure 1.**
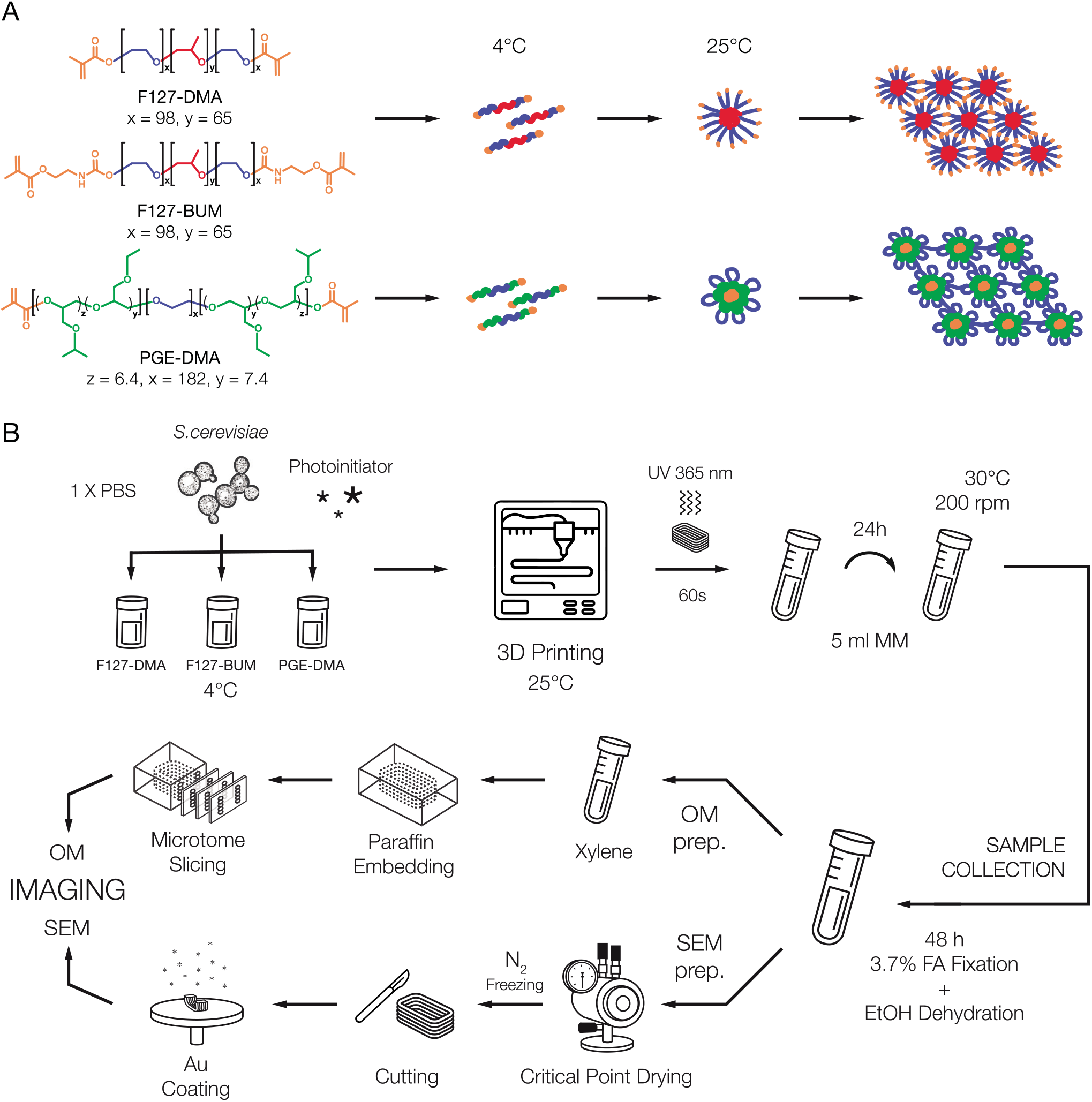
Schematic diagram showing polymer chemistry (A) and experiment workflow (B). The polymers were mixed with PBS, cells, and photoinitiator at 4 °C and printed at 25 °C to be cured after printing. Batch cultivation time was 24 h for varying days. Samples were collected, fixed, and dehydrated. Specific preparation protocols were applied for SEM or OM imaging.

## Results and Discussion

### Stability of cell-free hydrogels under physiological conditions

F127-DMA and F127-BUM hydrogels were prepared as 30 wt% in PBS buffer, while PGE-DMA was prepared as 20 wt%, and all formulations included 0.15 wt% 2-hydroxy-2-methylpropiophenone as a photoinitiator. These hydrogels have previously been shown to be printable using a direct-write extrusion printer.^4,17,18^ Despite the fact that F127-DMA and F127-BUM were present at the same concentration in their respective hydrogels, the latter polymer resulted in hydrogels that had a larger storage modulus (247 kPa versus 203 kPa).^4,19^ The data were acquired in milli-Q water, but the storage modulus pattern should remain relatively similar in PBS.^28^ The difference in stiffness of F127-based gels is attributed to the presence of carbamate linkage at the polymer chain ends in F127-BUM (Figure S1), which can form intermolecular hydrogen bonds. The PGE-DMA hydrogel had a lower storage modulus (96 kPa) largely due to the lower concentration of polymer present.^18^ Concentrations of PGE-DMA beyond 20 wt% were not possible as the hydrogel became too stiff for processing. The lower feasible concentration for PGE-DMA gel formation is attributed to the difference in the self-assembled networks. In particular, the presence of bridging chains in BAB triblock copolymer assemblies could facilitate the gelation (Figure 1A).

We first sought to understand how cells proliferate and affect the surrounding hydrogel matrix, which was observed using optical microscopy (OM) and scanning electron microscopy (SEM). The stability of the cross-linked hydrogels in the absence of any cells was observed for 14 d in a minimal medium (MM). The images presented here serve as a control (Figure 2) to appreciate the differences with yeast-laden hydrogels, where the structures might transform due to proliferation of cells. At both macroscopic and microscopic levels, the control samples appeared stable throughout the cultivation period under physiological conditions. Moreover, we did not observe any changes in the physiological environment as determined by glucose and pH measurements (Figure S2 A, B). The mass of the control structures remained unchanged.

**Figure 2.**
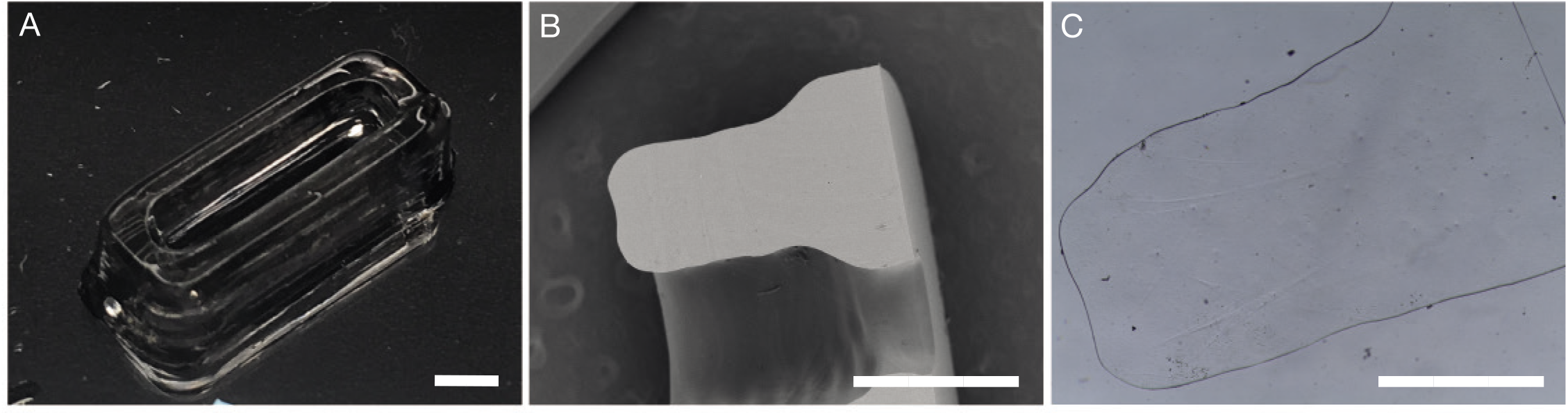
Illustrative images of control structures (hydrogels printed without cells). Photograph after 24 h equilibration (A). Cross-sectional SEM micrograph (B). OM micrograph, slice thickness 40 μm (C). Scale bar 1 mm.

### Stability of yeast-laden hydrogels

After ascertaining the stability of all hydrogels printed without cells, we focused on understanding the impact of long-term proliferation of immobilized cells on the hydrogels and whether different proliferation patterns were adopted by cells in the distinct LMs. Here, we printed the same formulations as mentioned before with a cell inoculum of 10^6^ cells g^−1^ hydrogel using a direct-write extrusion printer. After ultraviolet (UV) curing, we washed each LM for 60 s in 70 % ethanol to ensure sterility of printed structures and to avoid potential contamination of the culture medium from peripheral cells. We cultivated the LMs in 5 mL MM for 14 d in at least triplicates with a change of medium every 24 h. Representative samples were collected for processing either for OM or SEM on days 0, 7, and 14. During sample fixation and dehydration for microscopy analysis, some cells detached from the cross-section surface.

Starting on day 0 (after equilibration for 24 h at room temperature), small colonies were observed inside the 3D printed structures (Figure S4 A, C, E, G, I, K). After one-week, clear differences were observed in how cells grew inside each of the hydrogels; those distinct proliferation patterns remained largely consistent during the second week. Peripheral colonies in F127-based LMs tended to merge and formulate a separate film around LMs (Figure 3 A – B, D – E). These materials cracked open beyond a particular cell number (S4 D, J). For some samples in F127-DMA, the cell-free layer and cell-laden layer tended to separate completely (Figure S4 B, H; Figure S6). The growth of colonies in PGE-DMA was directed towards the periphery (Figure 3 C, F; Figure S4 F). A separated layer as in F127-based LMs was not observed. We observed that there was a colony diameter size gradient in all LMs, with smaller colonies in the middle and larger ones toward the periphery (Figure S5). Colony diameters in the middle of the structure for all three hydrogel compositions stayed in the range of 26-38 μm, with similar observations for day 7 and 14 samples (Figure 3; Figure S4). Cell-retaining structures became swollen due to cellular proliferation (Figure 3 G), and colonies in the middle region started to show an altered morphology indicating phenotypic differences in cells (Figure 3 I). Potentially, there was a limitation of nutrients for inner cells that contributed towards a clear colony size gradient (Figure S5); the cells in the smaller, nutrient-limited inner colonies were also likely more prone to cell death (Figure 3 I, S5). A similar pattern has been reported in a recent study by Qian and colleagues.^7^

**Figure 3.**
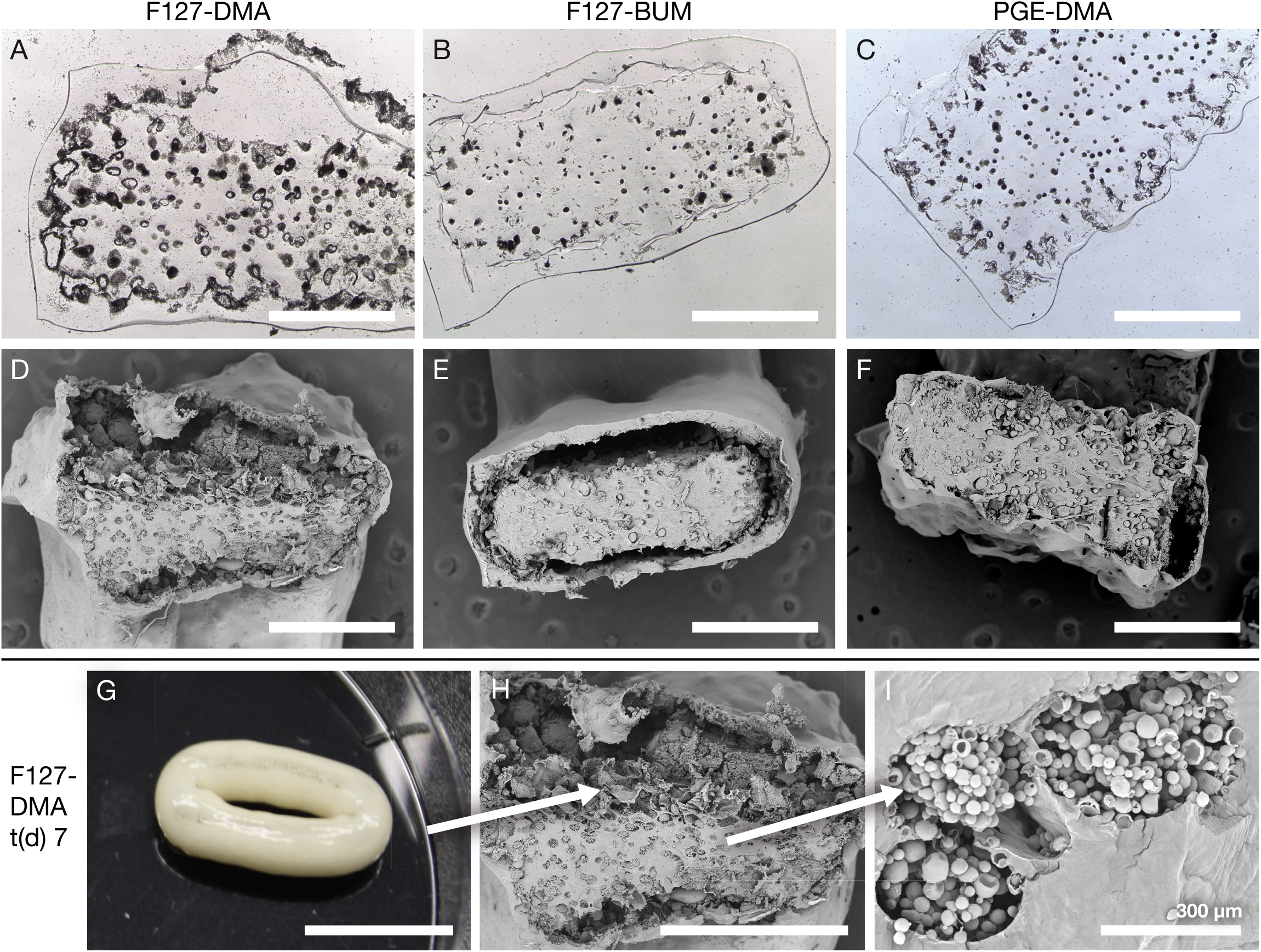
OM (A - C) and SEM (D - F, H, I) micrographs of LMs after 7 days of incubation. Cells escaped from PGE-DMA without major disruption of the material, F127-DMA and F127-BUM formed separated layers (center vs. shell). Most cell proliferation occurred at the interface. The LMs swelled up and retained cells up to a particular cell number (G). As peripheral colonies joined into one major colony (G, H) the colonies residing in the middle of the hydrogel were deprived of access to nutrients causing cell death in colonies (I). Scale bar 1 mm unless marked differently.

Two days after the start of the experiment, glucose was always depleted within every 24 h batch cultivation (Figure S2 C), and the pH did not drop substantially (Figure S2D). The cells started to escape into the culture medium at different time points. F127-DMA retained cells the longest (5.13 ± 1.55 d), whereas F127-BUM, although structurally almost identical, retained them for 3 d, and PGE-DMA for 2 d (Figure S2E). Due to growth, swelling, and retention time differences in F127-DMA and F127-BUM, the living materials were 136.80 (± 27.78) %, 53.39 (± 4.15) % respectively, heavier after one week, (Figure S2 F). The observed mass increase was only 41.28 (± 9.93) % for PGE-DMA (Figure S2 F). After the cells started escaping from hydrogels, they continuously washed out from the cavity between the inner and outer layer of F127-based materials and resulted in an increase of about 40 % (relative to the start) after two weeks of cultivation (Figure S2 F). However, since PGE-DMA did not form such cavities, its weight stayed the same during the second week (Figure S2 F). Interestingly, the increase in mass was the same (roughly 40 %) for all LMs after two weeks of incubation. This appears to be the carrying capacity of all tested living materials under our experimental conditions after two weeks and suggests that modifications to hydrogels would be necessary to increase the capacity in the future.

### Growth patterns of yeast colonies in living materials

To understand cellular growth within the hydrogels, we 3D printed these materials with a lower number of cells (~10^5^ g^−1^ hydrogel), allowing us to observe single colonies after 48 h of cultivation. Batch cultivation in 5 mL MM was carried out with a medium change every 24 h. As observed before, cells were retained in both F127-DMA and F127-BUM hydrogels on day two, whereas cells started to escape from PGE-DMA (Figure S7 A). Therefore, the comparison of glucose consumption in PGE-DMA hydrogels with F127-based hydrogels was not possible from the second day onwards. Within 24 h, glucose was consumed slightly faster in PGE-DMA and F127-DMA hydrogels compared to F127-BUM hydrogels, where the difference in starting/finishing glucose was only minimal (Figure S7 B). After 48 h, glucose was consumed significantly more in the medium of F127-DMA than F127-BUM, supporting the aforementioned observation (Figure S7 B). The ABA block architecture of F127-DMA and F127-BUM, and the BAB block architecture of PGE-DMA afford different physically and chemically cross-linked networks and storage moduli. Those differences, among others, further support differences in glucose diffusion through the hydrogel (Figure S7 B). Further studies are required to assess the diffusion of molecules through these hydrogel matrices, wherein the polymer composition, architecture, and concentration are altered in order to design or select other LMs based on these diffusion parameters.^29,30^

Interestingly, the morphology of colonies differed between F127-based LMs and PGE-DMA hydrogels. While the colonies in F127-based materials were spherical in shape (Figure 4 A – B, D – E), PGE-embedded colonies showed a more irregular spindle-like or elliptic shape (Figure 4 C, F, H). For this reason, it was impossible to properly measure and compare the colony size and growth rate inside PGE-DMA relative to the F127-based hydrogels. Additionally, lesions appeared on the surface of PGE-DMA hydrogels, confirming the escape of cells into the medium (Figure 4 I), which were not observed in F127-based hydrogels. After 72 h, the F127-based hydrogels had single colonies in the range of 90-250 μm, and the proliferation of the colonies toward the center of the hydrogel did not exhibit a significant change in size to the ones after 48 h (Figure S8).

**Figure 4.**
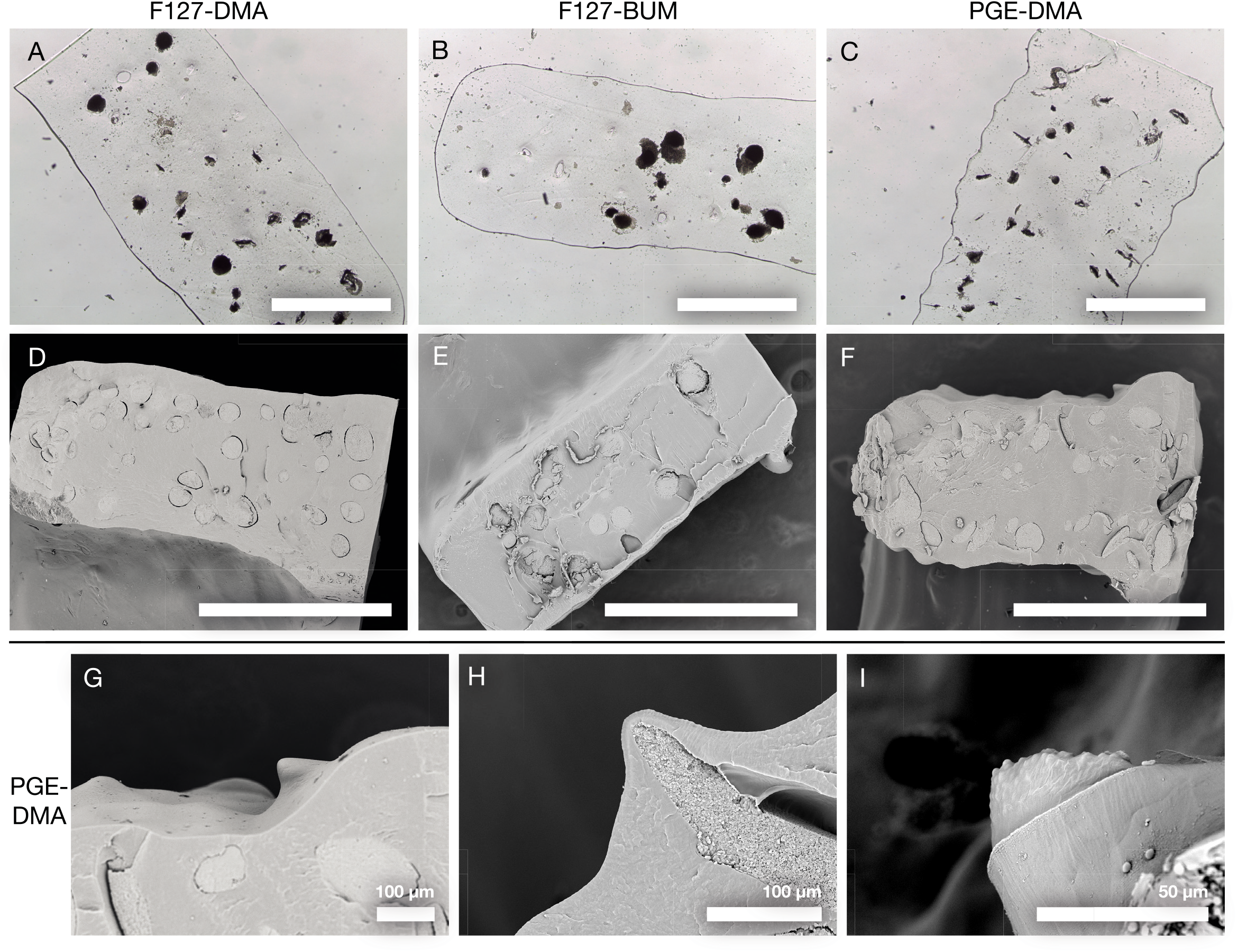
SEM and OM micrographs of LMs after 48 h of incubation (A - F) and the escape mode of cells from PGE-DMA (G - I). Peripheral colonies in PGE-DMA formed spindle-like structures (C, F, H), while F127-DMA and F127-BUM formed spherical colonies (A, B, D, E). When the material broke, cells started escaping into the medium (I). Scale bar 1 mm unless marked differently.

### Cellular phenotyping in living materials

Further investigations of the cell-laden hydrogels revealed a thin organic coating around the cell colonies in the F127-DMA and F127-BUM hydrogels (Figure 5 A, B), which was not present in the PGE-DMA hydrogels (Figure 5 C). This difference became evident within 24 h after 3D printing and incubation at room temperature. A supercritical CO_2_ extraction protocol was used with rapid release of CO_2_ to separate and measure the thin polymer coating (100 – 160 nm) around yeast colonies (Supplement 3.2). As colonies increased in size, the thin surrounding coating ruptured, and only remnants of the coating were observed on a colony surface (Figure S9). Based on SEM images, we determined that the film ruptured when the colony diameter had reached a size of about 60 – 80 μm within the LMs (Figure S9).

**Figure 5.**
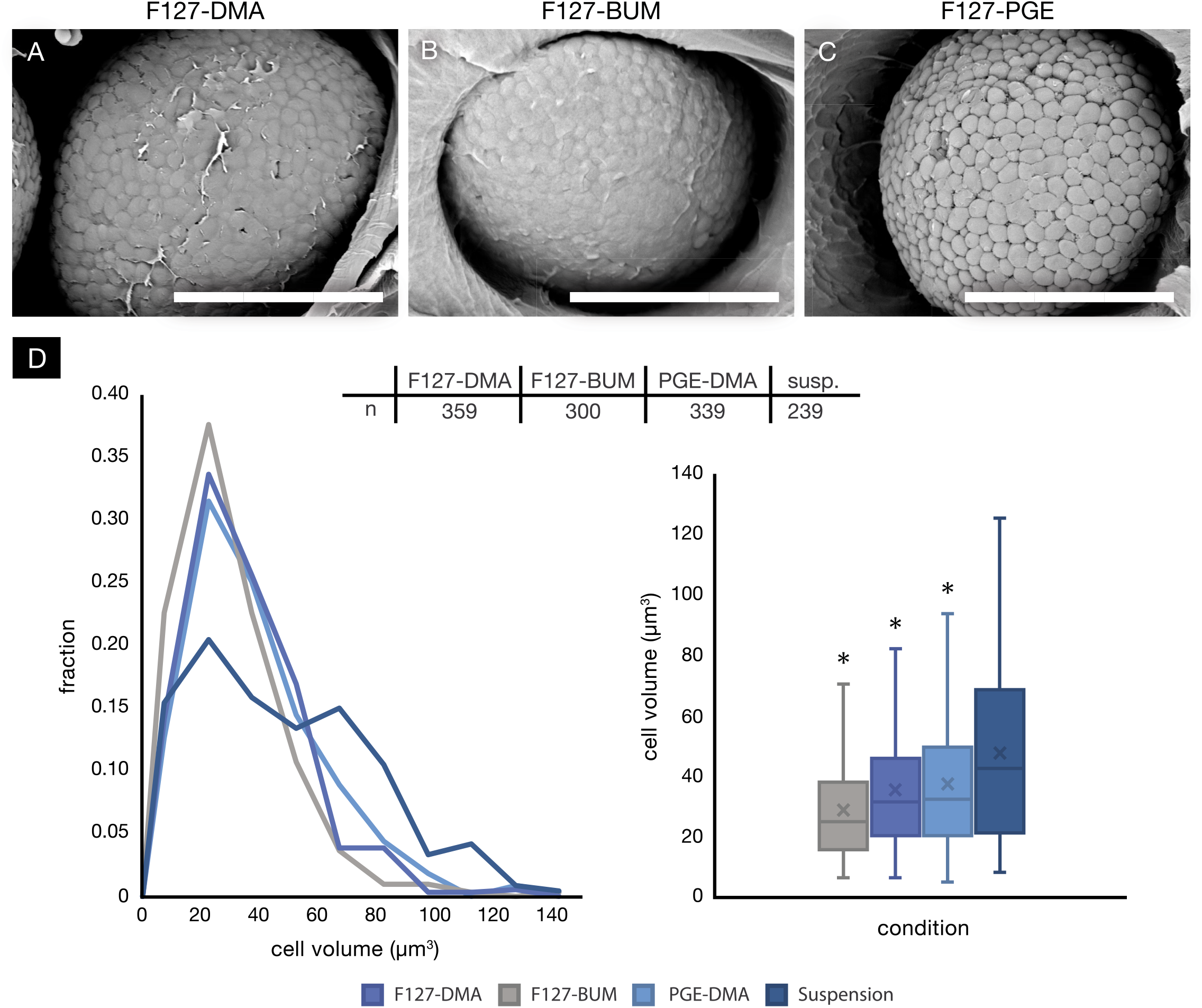
Organic film covers yeast cell colonies in LMs and cell size (μm^3^). F127-DMA (A) and F127-BUM (B) both had a thin organic coating around colonies, while PGE-DMA (C) lacked a similar polymer coating. Scale bar 20 μm. Size (in μm^3^) distribution of cells in LMs and suspension cell culture (D). A total of ≥ 239 cells per condition were analyzed. LMs were incubated for 48 h before processing and the measurement. * *p* < 0.05, significant difference from suspension cells, x: mean.

The different cell-polymer interactions, as well as retention times, consequently led to the question of impact of physical confinement on the cell phenotype. To address this question, we performed a computational analysis on the acquired SEM images as described in the materials and methods section. During this analysis, cell size (in μm^3^) parameter was utilized to investigate the effects of physical confinement on the cells in hydrogels in comparison to suspension cells. We evaluated cell size differences after 48 h in the aforementioned samples (used in Figure 4). Suspension cells were cultivated, fixed and dehydrated in the same way as the immobilized cells (Figure S10). Interestingly, cells encapsulated in hydrogels were significantly (*p* < 0.05) smaller than the suspension cells (Figure 5D). Cell size differences were even evident among the LMs (Figure 5D). The cause of cell size differences was likely multifarious instead of an individual attributable factor, as a cellular phenotype is an integrated readout of manifold cellular processes and naturally, physical confinement in hydrogels is an additional factor for immobilized cells compared to suspension cells (Figure 5). The previous studies on effects of physical confinement in a calcium alginate matrix indicate changes in cellular physiology of yeast.^31,32^ Though, a molecular investigation of cells was not in the scope of the present study, it would be very valuable to understand underlying molecular mechanisms responsible for phenotypic differences in LMs for their development as a technology of the future.

### Polymer integrity analysis in living materials

In certain applications polymer integrity could be of essence in the LMs but in other instances a controlled polymer degradation might be preferred, making the study of polymer degradation an important component for development of the LM based technologies. F127-BUM contains carbamate bonds on the periphery of micelles making it susceptible to enzymatic degradation by cells, which can secrete proteases^32^, and thus provides an excellent model for studying polymer degradation (Figure 1; Figure 6). To validate this idea, we conducted a 14-day experiment with and without a protease inhibitor cocktail for all the LMs in the study (Figure 6). The concentration of inhibitors used in these experiments did not have an influence on control structures nor on the cell proliferation. We used the fast release of gas in the supercritical CO_2_ extraction protocol to identify differences between degraded and intact polymers in LMs (Figure 6, Supplement 3.2).

**Figure 6:**
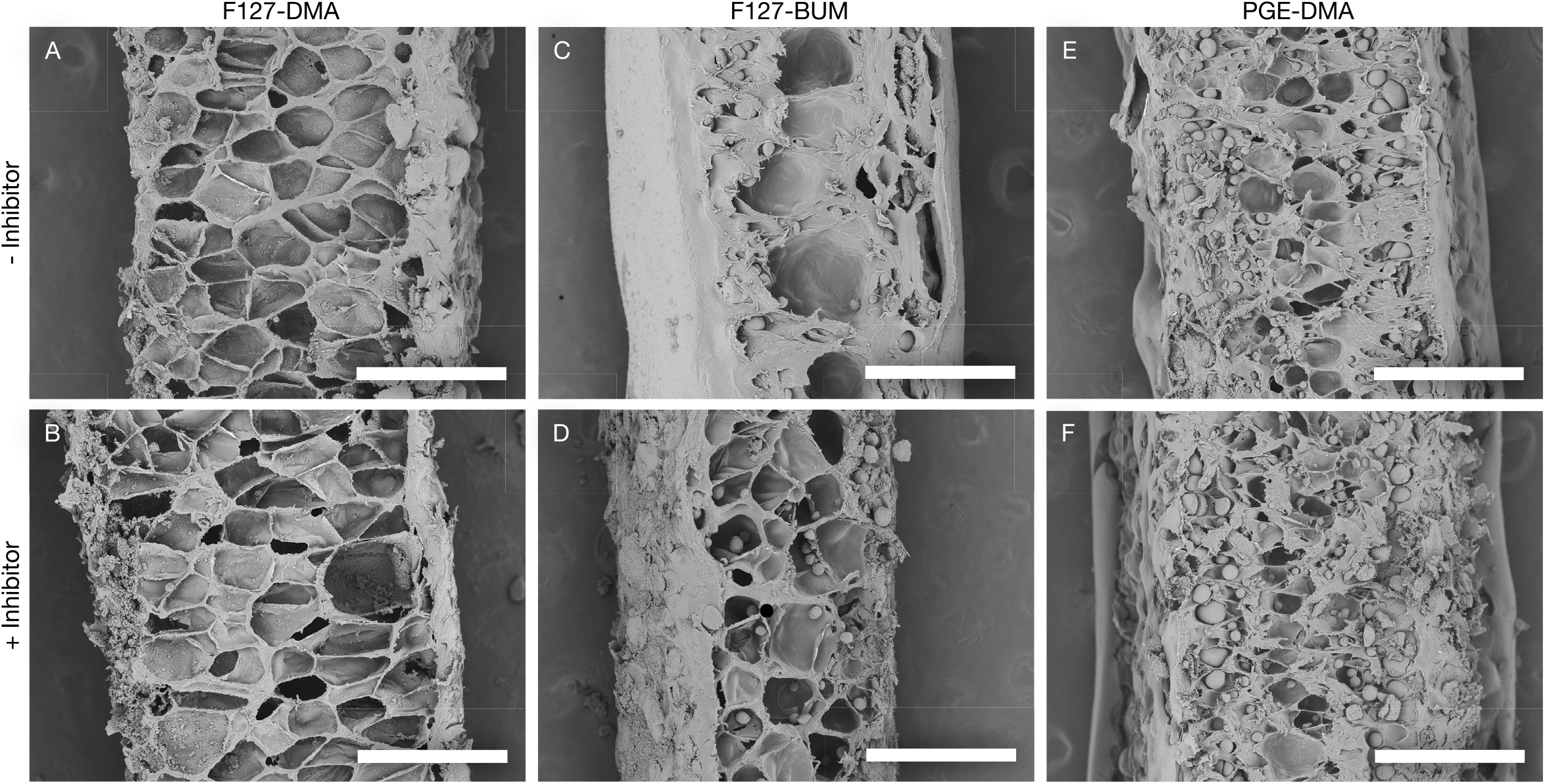
SEM micrographs showing cavities after pressure release of CO_2_ inside LMs. Upper panel: normal cultivation; lower panel: enzyme inhibitors added into the MM. No visual differences were detected in DMA-functionalized polymers (A – B, E - F). Differences were observed in F127-BUM (C, D): when the enzyme inhibitors were missing, the material was degraded allowing the gas to escape more easily during sample preparation (C), whereas gas got trapped in intact material and formed more cavities (D). 24 h 5 mL batches over 14 days. Scale bar 500 μm. All samples prepared in the single critical CO_2_ extraction experiments to avoid technical variations.

The addition of inhibitors, as expected, did not have any effect on the outcome with F127-DMA and PGE-DMA compared to the control condition - shown by similar images (Figure 6 A – B, E – F). It should also be noted that in the case of PGE-DMA, the effect of the fast gas release appeared less pronounced compared to F127-based materials. This result could be attributed to the lower polymer concentration used in the formation of PGE-DMA hydrogels. During fast CO_2_ extraction, less dense (or degraded) materials allow the gas to escape more easily (Figure 6 C, E, F), whereas denser materials withhold the gas resulting in cavities (Figure 6 A, B, D). A clear difference was observed in the F127-BUM samples, where the sample appeared altered in the absence of the inhibitor indicating a polymer degradation (Figure 6 C, D). To ensure the reproducibility of these observations, we repeated the experiment for F127-BUM with a culture medium change every 48 h allowing secreted proteases more reaction time to potentially cleave the bonds. Following this approach, we found an even more pronounced difference, indicating a possible effect of enzymes on the integrity of F127-BUM (Figure S3.2 E, F). This was also supported by a higher cell retention time in F127-DMA compared with F127-BUM (Figure S2 E). Using different proteases to study the enzyme degradation of F127-BUM might be worth an investigation in the future, and can potentially make it an attractive candidate for use in biomedical applications, such as angiogenesis research, where enzyme-driven matrix degradation is vital.^33^

## Conclusion

Our ability to develop new polymeric materials and their hydrogels for 3D printing LMs is outpacing our understanding of LMs due to the lack of investigations into how the mutual interactions of incorporated cells in the living materials impact both the cells and the polymers. Understanding such cellular-polymeric interactions is crucial to draw conclusions about the effects of physical confinement on cells within these materials.

We investigated three triblock polymeric hydrogels together with yeast-laden LMs with scanning electron and optical microscopy techniques and revealed several previously unreported features, *e.g*. proliferation patterns of colonies, alteration of cell sizes, colony encapsulations, and potential polymer degradation. The distinct compositional and structural features of the living materials make them suitable for different applications, *e.g*. in biomedical research, where degradable polymers are needed, or biotechnology, where cell retention is desirable. Factors, such as the printing thickness and diffusion, should also be considered to ensure a sufficient nutrient supply to all cells within the LMs. Here, we have demonstrated changes in cellular phenotypes due to physical confinement within three hydrogels. However, our current study precludes understanding underlying molecular mechanisms of phenotypic changes, but it would constitute an important area of exploration in the future.

## Materials and Methods

### Chemical Synthesis of Polymers

Two F127-derived polymers, namely F127-BUM & F127-DMA, and PGE-DMA were provided by the Nelson laboratory at the University of Washington. Synthesis as mentioned in the laboratory’s papers by Saha *et al*. (2018), Millik *et al*. (2018) and Johnston *et al*. (2019). The percent (%) functionalization for F127-DMA was 81 %, while F127-BUM and PGE-DMA were functionalized quantitatively.

#### F127-DMA

The F127-DMA polymer used in this study was synthesized using methacryloyl chloride as described in the literature.^4^ Briefly, Pluronic F-127 was dried and subsequently dissolved in anhydrous toluene under a nitrogen atmosphere. Triethylamine was added and the solution was cooled to 0 °C. A solution of methacryloyl chloride in anhydrous toluene was added dropwise to the solution. After complete addition of the methacryloyl chloride solution, the reaction mixture was warmed to room temperature and allowed to stir for 24 h. The polymer was collected via vacuum filtration, concentrated under reduced pressure, and reconstituted in fresh toluene. This process was repeated two more times. The polymer was once again dissolved in toluene and precipitated in diethyl ether. The polymer was rinsed twice with fresh ether and collected via centrifuge. The polymer was dried in a vacuum oven to afford a fluffy, white powder.

#### F127-BUM

The F127-BUM polymer used in this study was synthesized using 2-isocyanatoethyl methacrylate and dibutyltin dilaurate according to the literature.^19^ Briefly, Pluronic F127 was dried and subsequently dissolved in anhydrous dichloromethane (DCM). Dibutyltin dilaurate was added to the mixture, followed by the dropwise addition of a 2-isocyanatoethyl methacrylate/DCM solution. The reaction was allowed to stir for 2 d, quenched with methanol, and precipitated in diethyl ether. The polymer was collected via centrifuge and washed twice with fresh ether. The polymer was dried under vacuum to afford a fluffy, white powder.

#### Unfunctionalized PGE

The unfunctionalized PGE precursor polymer was synthesized by anionic ring opening polymerization as described in the literature.^11^ Briefly, PEO was added to the reaction vessel and dried under vacuum overnight. Dry THF was added and a potassium naphthalenide solution was titrated into the flask under an argon atmosphere. Isopropyl glycidyl ether and ethyl glycidyl ether were added simultaneously to begin polymerization. The reaction continued for 24 h at 65 °C. The reaction mixture was then precipitated into cold hexane and washed twice. The isolated polymer was dried in a vacuum oven to afford the unfunctionalized PGE polymer precursor as an off-white solid.

#### PGE Methacrylate

The methacrylate functionalized PGE polymer used in this study was synthesized using methacrylic anhydride as described in the literature.^18^ Briefly, the PGE polymer precursor was dissolved in dry THF under a nitrogen atmosphere. Triethylamine was added and the mixture was heated at 65 °C for 30 min. Methacrylic anhydride was then added and the mixture was stirred for 16 h at 65 °C. The reaction was then precipitated into cold ether. The polymer was collected and washed twice with additional ether, once with hexane, and dried under vacuum for 24 h to afford the methacrylate functionalized PGE polymer (PGE-DMA) as an off-white solid.

### Yeast Strain, Media and Cultivation Conditions

The yeast strain *S. cerevisiae* CEN.PK113-7D (*MATa, MAL2-8^c^, SUC2*) was used throughout the study and cultivated in MM. The composition of 1 L MM (pH = 6.9) was 10 g glucose (Acros Organics), 2.5 g of (NH_4_)_2_SO_4_ (Lach-Ner), 3 g of KH2PO4 (Sigma-Aldrich), 5.25 g of K_2_HPO_4_ (Merck) and 0.25 g of MgSO_4_ (Sigma-Aldrich) in milli-Q water. One milliliter trace elements (all Sigma-Aldrich, unless marked differently) and 1 mL vitamin solution (all Sigma-Aldrich, unless marked differently) were added after sterilization of the MM. One liter trace element solution (pH = 4) contained EDTA sodium salt (Lach-Ner), 15.0 g; ZnSO_4_·7 H_2_O, 4.5 g; MnCl_2_·2 H_2_O, 0.84 g; CoCl_2_·6 H_2_O, 0.3 g; CuSO_4_·5 H_2_O, 0.3 g; Na_2_MoO_4_·2 H_2_O, 0.4 g; CaCl_2_·2 H_2_O (Carl Roth), 4.5 g; FeSO_4_·7 H_2_O, 3.0 g; H3BO3, 1.0 g; and KI, 0.10 g. One liter vitamin solution (pH = 6.5) contained biotin, 0.05 g; p-amino benzoic acid, 0.2 g; nicotinic acid, 1 g; Ca pantothenate, 1 g; pyridoxine-HCl, 1 g; thiamine-HCl, 1 g; and myoinositol (AppliChem), 25 g; in milli-Q water. Where indicated, a SIGMAFAST inhibitor cocktail (S8820, Sigma Aldrich) was used in MM at a concentration of 0.1x according to the manufacturer.

Cell cultivation was carried out in 15 mL tubes (5 mL MM) at 30 °C and 200 rpm in an incubator. The living materials were washed in 70 % ethanol (Berner Pro) for 60 s after printing and equilibration to avoid contamination and viable yeast on the surface of living materials. A short wash in 70 % ethanol was applied after every 24 h batch.

### Hydrogel Preparation for 3D Printing

Sterile phosphate-buffered saline (PBS) solution was mixed with a desired polymer and cooled at 4 °C overnight to prepare a hydrogel (F127 hydrogels: 30 wt%, PGE-DMA: 20 wt%). One liter of PBS (pH = 7.2) contained 8 g of NaCl (Sigma-Aldrich), 1.44 g of Na2HPO4 (Fisher Scientific), 0.24 g of KH2PO4 and 0.2 g of KCl in milli-Q water. To make a hydrogel ready for printing, 1.5 μL g^−1^ hydrogel of the photoinitiator 2-hydroxy-2-methyl propiophenone (Irgacure 1173; >97 %, Sigma-Aldrich) were mixed in at a temperature of 4 °C. If needed, 10^5^ or 10^6^ spun-down cells g^−1^ hydrogel were added. A short stirring of both additives ensured an equal distribution and after incubating 30 min on ice, to make the solution bubble-free, it was poured into a 10 mL dispensing barrel equipped with a 0.41 mm dispensing tip (both Adhesive Dispensing) and warmed to room temperature to transform to a shear-responsive state for printing.

### 3D Printing

3D printing was performed on a K8200 printer (Velleman, Belgium) modified to be applicable for direct pressure dispensation. The computer-aided design model was designed with Solidworks (Student Edition) and the G-code was generated using open source 3D printing toolbox (Slic3r 1.3.0). The model’s measures were 10 x 3 x 3.5 mm (X, Y, Z) sliced with one outer perimeter and printed in vase mode with print speed of 10 mm s^−1^. Directly after the print, the hydrogel was cross-linked for 60s with four light-emitting diodes (CUN66A1B, Seoul Viosys, Republic of Korea) emitting at a wavelength of 365-367 nm.

### High-Performance Liquid Chromatography

Chromatography was performed using an Aminex HPX-87H Column (Bio-Rad, USA) with 5 mM sulfuric acid (>99.5 %, Merck) as a mobile phase at 45 °C. A Shimadzu Prominence-i LC-2030C Plus (Japan) equipped with a Refractive Index Detector RID-20A (Shimadzu, Japan) was used to detect the components.

### Fourier-Transform Infrared Spectroscopy

FTIR spectroscopy was used to obtain structural information about the polymers. Polymers were dried for 24 h at 25 °C in 1 mbar vacuum (VO200, Memmert, Germany). The measurements were performed using an Alpha spectrometer equipped with Platinum ATR (Bruker, USA). The polymers were analyzed over the range of 3800–400 cm-1 and averaging was over 24 spectra each.

### Macroscopic Observations

Samples were arranged and imaged on a petri dish after the indicated amount of time. Pictures were acquired with a Canon EOS 450D equipped with a Canon Zoom Lens EF 17 – 40 mm.

### Scanning Electron Microscopy

a. Sample Fixation and Dehydration The samples were fixed for 48 h in 3.7 % formaldehyde (Biotop/Naxo) in 0.1 M phosphate buffer (PB) fixation solution, which was replaced after 24 h. One liter of 0.2 M phosphate buffer contained 20.44 g of Na2HPO4 and 6.72 g of NaH2PO4 (Acros Organics). For sample dehydration, 99.5 % ethanol was used to establish several dilutions of it in milli-Q water. Samples were dehydrated in an ascending ethanol series (40 – 90 %, 10 % steps; 96 %, 99.5 %) at room temperature (2 h minimum per step, last step overnight followed by replacement with fresh absolute ethanol).
b. Critical Point Drying Critical point dryer (E3100, Quorum Technologies, United Kingdom) was cooled to 15 °C with a thermostat (Proline RP 1845, LAUDA, Germany). The samples were mounted on a tray and inserted into the critical point dryer. The dryer was filled with liquid CO_2_ to replace the ethanol and the chamber was purged 6 – 8 times in 30 – 60 min intervals. The critical point was reached by increasing the temperature to 37 °C and controlling the pressure not to exceed 110 bar, followed by pressure release to recover the samples. The pressure release was done either fast or slow. With slow release, pressure was released slowly overnight until the chamber was ready to be opened. With fast release, pressure from 110 to 60 bar was released slowly (to avoid cooling the reactor and turning supercritical state back to liquid state) and from there, the remaining pressure was released within 3 min. The samples shrank by 35 – 40 % due to the drying process.
c. Sample cutting The samples were frozen in liquid N2 and cut with a scalpel. For acquiring artifact-free cross-sections, the sample and scalpel, where immersed in liquid N2 for 20 s and instantly cut with fast incisions. For acquiring information of colony-material interactions, colony size and shape, the sample and scalpel where immersed in liquid N2 for 10 seconds and then cut after 2 – 3 s at room temperature with slow incision.
d. Sputter Coating A sample stub was covered with sticky carbon tape and the cut sample was attached on it. The sample was coated with a 7.5 nm thick gold layer using a high vacuum sputter coater (EM ACE600, Leica Microsystems, Germany).
e. Imaging Gold-coated samples were imaged with a tabletop scanning electron microscope (TM-3000, Hitachi, Japan), back-scattered electron detector. The imaging was done under a high vacuum and 15 kV accelerating voltage. Results were confirmed by imaging several samples over multiple slices. Colony sizes were directly detected using the measurement tool of the imaging software.
f. Image analysis SEM image analysis was performed using MATLAB2019b with Image processing and Parallel processing toolboxes. In this analysis, for cell size (μm^3^) calculations, yeast cells were assumed to have ellipsoid shape. The analysis relied on border detection using gradient magnitude between adjacent pixels. Different levels of smoothing and thresholding were applied to determine initial borders. Subsequently, watershed segmentation was applied and followed by pruning of borders using smoothing and thresholding techniques. The resulting borders were optimized using machine learning to reach the maximum possible score for each pixel, based on gradient intensity. The resulting borders were then used to segment the cells. Segmentation mask was then refined to remove parts with too high gradient intensity. The resulting cells were then filtered to remove any incorrectly segmented cells based on a score assigned to each segmented cell. Score was comprised of minimum and mean gradient intensities of a cell border, ratio of major and minor axes, solidity and cell area.

### Optical Microscopy

a. Sample Preparation The sample fixation was carried out in the same way as mentioned for SEM and finally transferred to histo-grade xylene (J.T. Baker) for 1 h. The samples were then placed into paraffin-embedding cassettes and covered with liquid paraffin (Leica) at 65 °C for 1 h to ensure proper infusion. The sample was taken out, orientated on a metal tray and covered with liquid paraffin. The sample was then cooled down.
b. Sectioning & Rehydration For microtome sections, a paraplast-embedded sample was mounted onto the microtome (RM2255, Leica Microsystems, Germany) and several slices were cut each. The slice thickness was 40 μm. Slices were collected from a water bath (milli-Q water) on a glass slide. The samples were dried overnight and then sequentially rehydrated in histo-grade xylene (20 min), 99.5 % ethanol (20 min), 90 % ethanol (20 min) and finally dH2O (20 min).
c. Imaging The rehydrated slices were carefully mounted on microscope glass slides and covered with ca. 40 μl of milli-Q water and a cover glass. A DM750 microscope equipped with an ICC50 HD camera system (both Leica Microsystems, Germany) was used. Results were confirmed by imaging several slices.

## Acknowledgements

This project has received funding from the European Union’s Horizon 2020 research and innovation program under grant agreement No. 668997, and the Estonian Research Council (grant PUT1488P). TB and HP would additionally like to acknowledge mobility grants MB-2018-1/25 by *Bayerisches Hochschulzentrum für Mittel-, Ost-und Südosteuropa* (BAYHOST) and the European Regional Development Fund, respectively. AN acknowledges the National Science Foundation (Grant No. 1752972), UW CoMotion Innovation Grant and UW Royalty Research Fund for financial support of this work. We thank Külli Jaako and Monika Jürgenson, at the Institute of Biomedicine and Translational Medicine, Faculty of Medicine, University of Tartu, for access to microscopy resources.

## Conflict of Interest

None

## Supplemental captions

**Figure S1: FTIR spectra of the three polymers**. The FTIR spectra of all 3 polymers appeared largely similar but F127-BUM clearly showed carbamate functional groups (–CON–) at 1532 cm-1. The spectrum also had less transmittance at 1720 cm-1, which validates one extra carbonyl (C=O) bond per repeating unit as compared to F127-DMA and PGE-DMA. Spectral lines of PGE-DMA and F127-DMA overlap.

**Figure S2: Medium/LM analysis upon cultivation over different times.** Control structures (A, B). LMs or their respective medium (C - F). MM glucose concentration over time (A, C). Average pH after each batch (B, D). Cell retention time (E). LM mass differences from starting day to day 7 or 14 (F). Gluc: glucose. * *p* < 0.05, significant difference from F127-DMA.

**Figure S3.1: Freeze-drying vs supercritical CO_2_ extraction**. A (simpler) alternative to the process of supercritical CO_2_ extraction is freeze-drying; their performance is compared by presenting an F127-BUM freeze-dried sample (A) as compared to a supercritical CO_2_ extracted one (B). A closer look at surface topology of the freeze-dried sample (C) shows a “brain”-like surface finish caused by uneven drying, in sharp contrast to the supercritical CO_2_ extracted sample (D). Freeze-drying can be used as an alternative for studying cross-sections of these materials, but with the caveat that the samples need to be sectioned after the drying process, because of ~20 μm deformations in the outer perimeter (E) as opposed to the supercritical CO_2_ extracted sample (F). Here is an overview of a freeze-dried sample (A) compared to a supercritical CO_2_ extracted sample (B). The surface topology of the freeze-dried sample (C) and the supercritical CO_2_ extracted sample (D). Cross-section and perimeter comparison of freeze-dried (E) and supercritical CO_2_ extracted (F) sample. F127-BUM samples displayed.

**Figure S3.2: Special features of fast pressure release during CO_2_ extraction**

Fast CO_2_ release at the end of supercritical CO_2_ extraction process generates local stresses in the gel that lead to mechanical changes resulting in a foam-like structure (A) and the separation of regions in the material (B, C) based on the local structural and mechanical properties. The amount and size of the generated pores depends on the speed of gas release. This method is a valuable tool for studying gas retention in the material, the thickness of thin film coatings around colonies (D) and possible degradation of the matrix (E, F). Foam-like structure generated due to fast CO_2_ release at the end of supercritical CO_2_ extraction (A). Separation (B, C) and measurement (D) of the thin polymer coating around yeast colonies in F127-BUM after 48 h of incubation. F127-BUM samples with (F) and without (E) protease inhibitor showing that when inhibitor was missing, the gas did not get trapped with fast CO_2_ release (pictures taken after 14 d with 48 h media change).

**Figure S3.3: Sample cutting - exposing cell-material interactions in hydrogels.** Varying the sample and the blade temperature together with the speed of cutting can be used to demonstrate various aspects of LMs. A combination of sample and scalpel cooling (~20 s) together with fast incisions results in the most accurate SEM images in terms of polymeric material and colony localization (A, B), but with this technique it is impossible to evaluate the colony size and shape because of the unknown location of the obtained cross-section in respect to the colony. A shorter duration of sample and scalpel cooling (~10 s) together with slow incision highlights biologically relevant information such as cell-polymer encapsulations (Figure 5 A - C) and colony size and shape (C, D) but results in cutting marks across the polymer (D). Different sample cuttings and resulting images: samples prepared with longer cooling of sample and scalpel showing relatively smooth cuts (A, B). Samples prepared with short sample and scalpel cooling showing clear colonies (C, D).

**Figure S4: OM and SEM Micrographs of LMs after 14-day cultivation:** OM (A-F) and SEM (G-L) micrographs of LMs after 0 (A, C, E, G, I, K) and 14 (B, D, F, H, J, L) d of incubation. Small colonies can already be observed on day 0. Disrupted LMs on day 14 with full separation of layers in F127-DMA (B, H). Scale bar 1 mm.

**Figure S5. Cell colony size gradient.** OM micrograph of PGE-DMA, day 14, slice thickness 150 μm (A). SEM micrograph of PGE-DMA, day 14 (B). Scale bar 500 μm. PGE-DMA illustrative for all LMs.

**Figure S6: Separation of outer layer**. Photo of F127-DMA after 14 d of incubation. Full separation of inner (black arrow) and outer (grey arrow) layer. Scale bar: 10 mm.

**Figure S7: OD and glucose consumption after 48h.** OD of MM after 48 h (A). Glucose concentration of MM with different LMs (B). * *p* < 0.05, significant difference from F127-DMA.

**Figure S8: SEM micrographs of LMs after 72 h incubation.** F127-DMA (A), F127-BUM (B), PGE-DMA (C).

**Figure S9: Polymer coating remnants on colony surfaces.** Remnants started appearing after the colony reached a certain size (60-80μm) in F127-based materials.

**Figure S10: SEM micrograph of yeast suspension cells.**

