## Supplementary figures and images for "Physical confinement impacts cellular phenotype within living materials"

### Supplemental S1

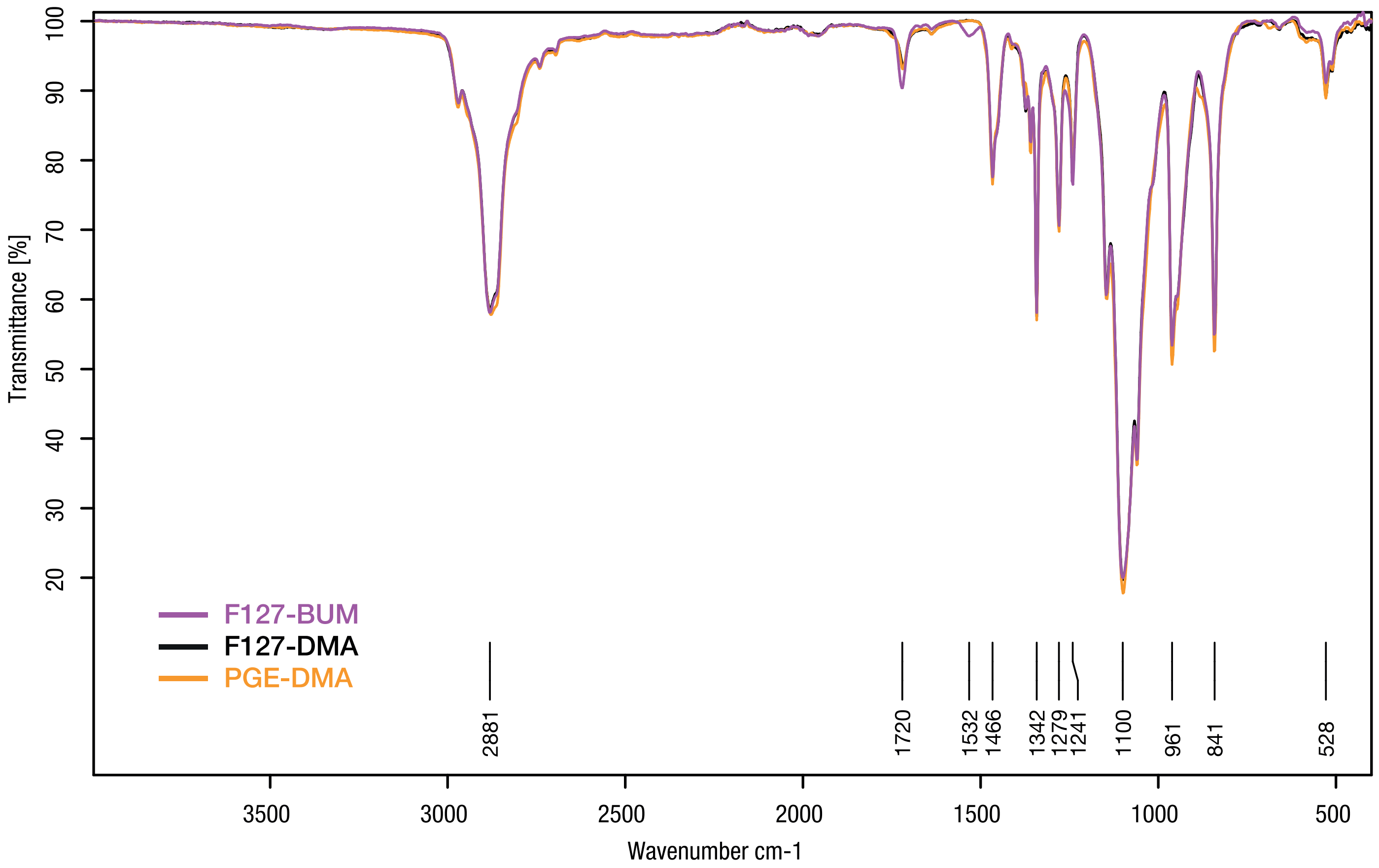

### Supplemental S3.1

Figure S3.1

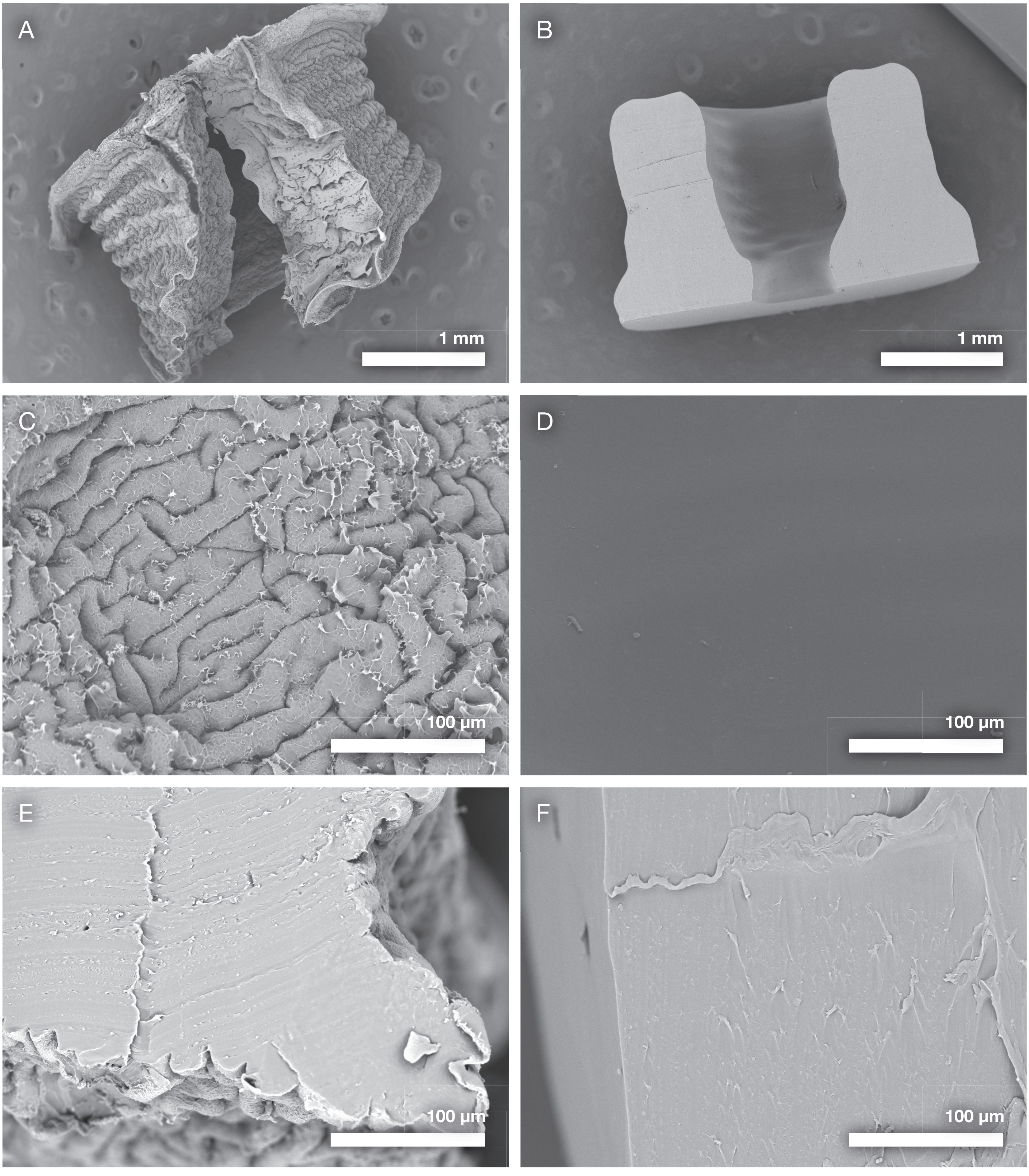

### Supplemental S3.2

Figure S3.2

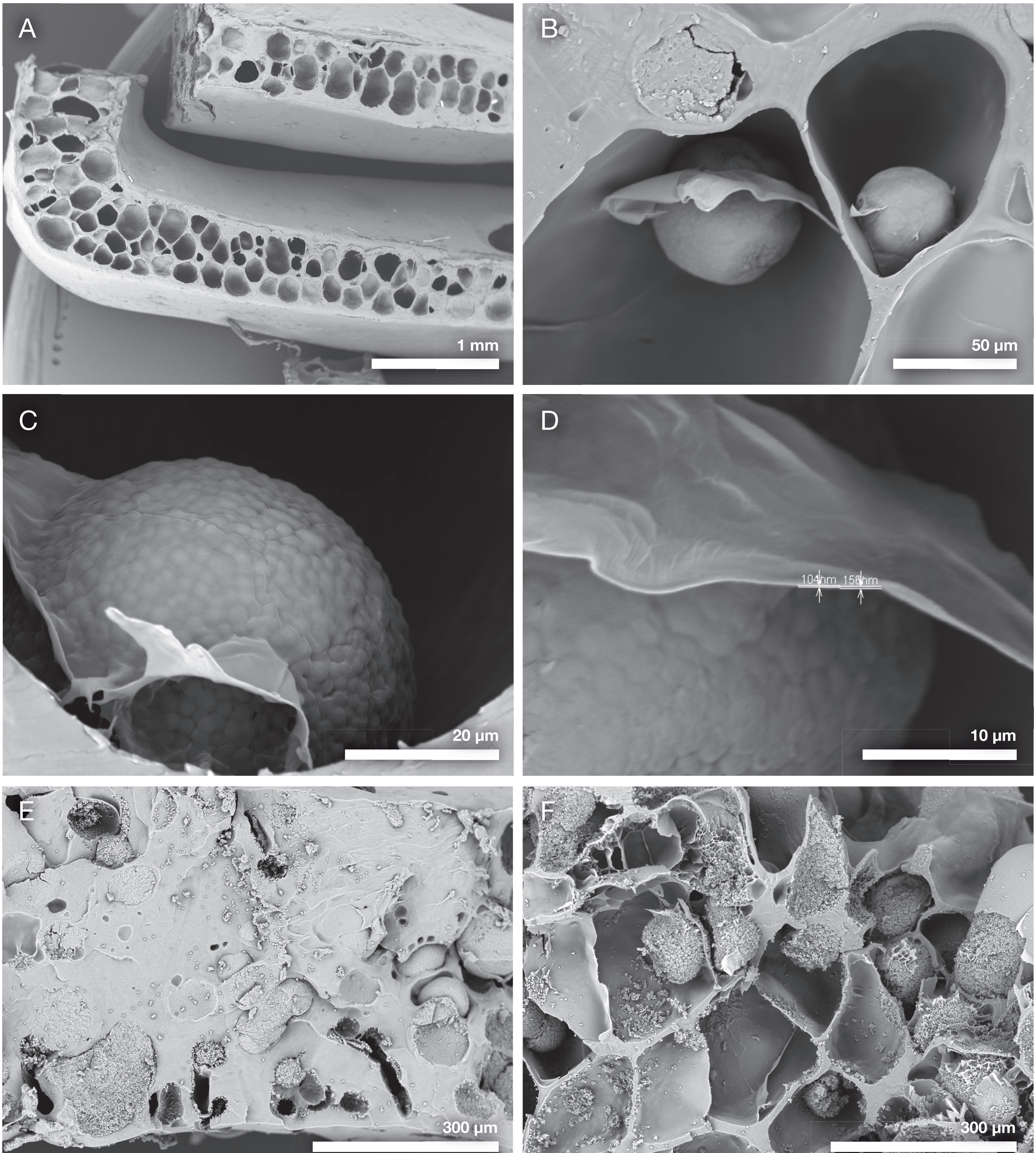

### Supplemental S3.3

Figure S3.3

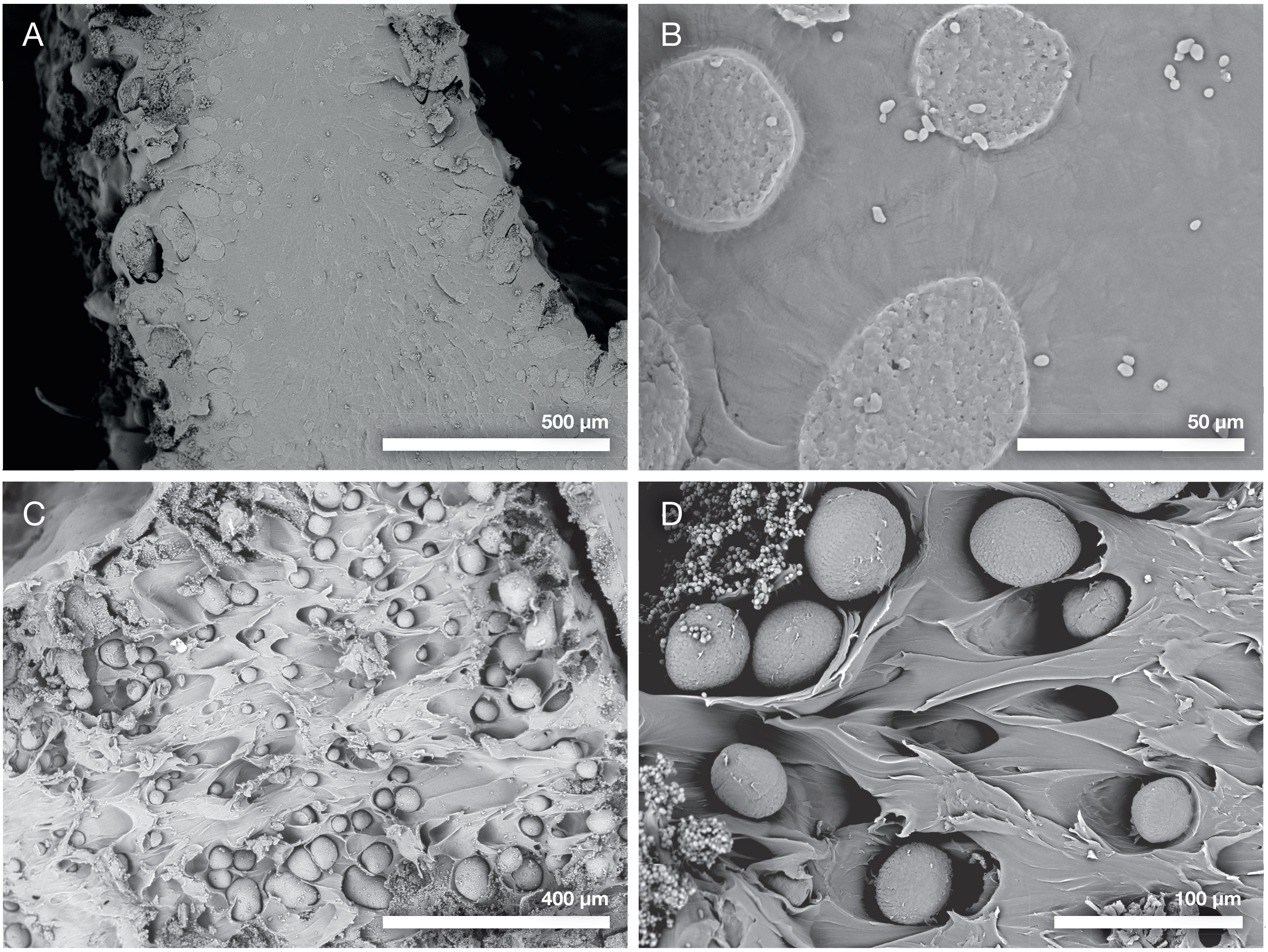

### Supplemental S4

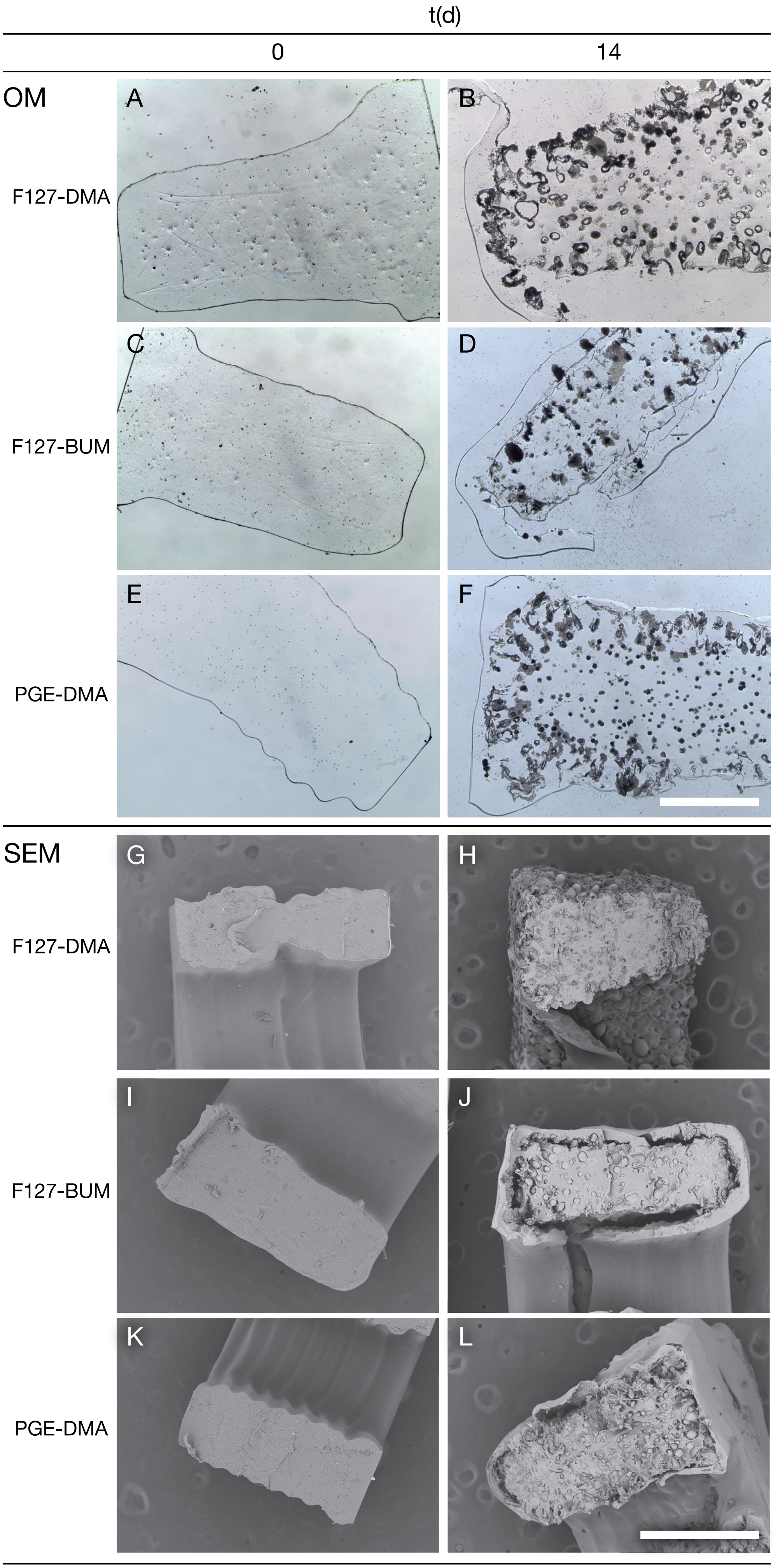

### Supplemental S5

Figure S5

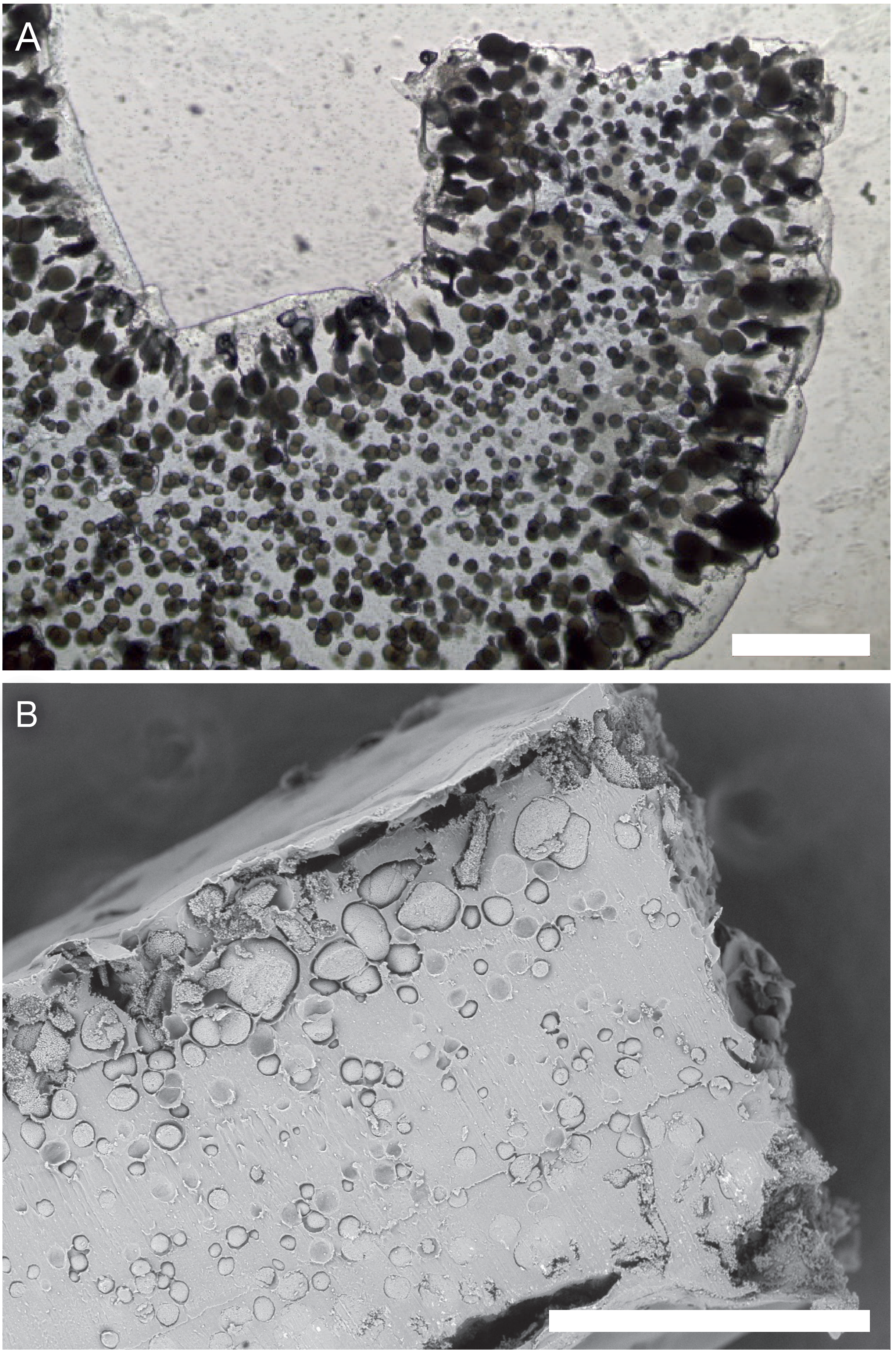

### Supplemental S6

Figure S6

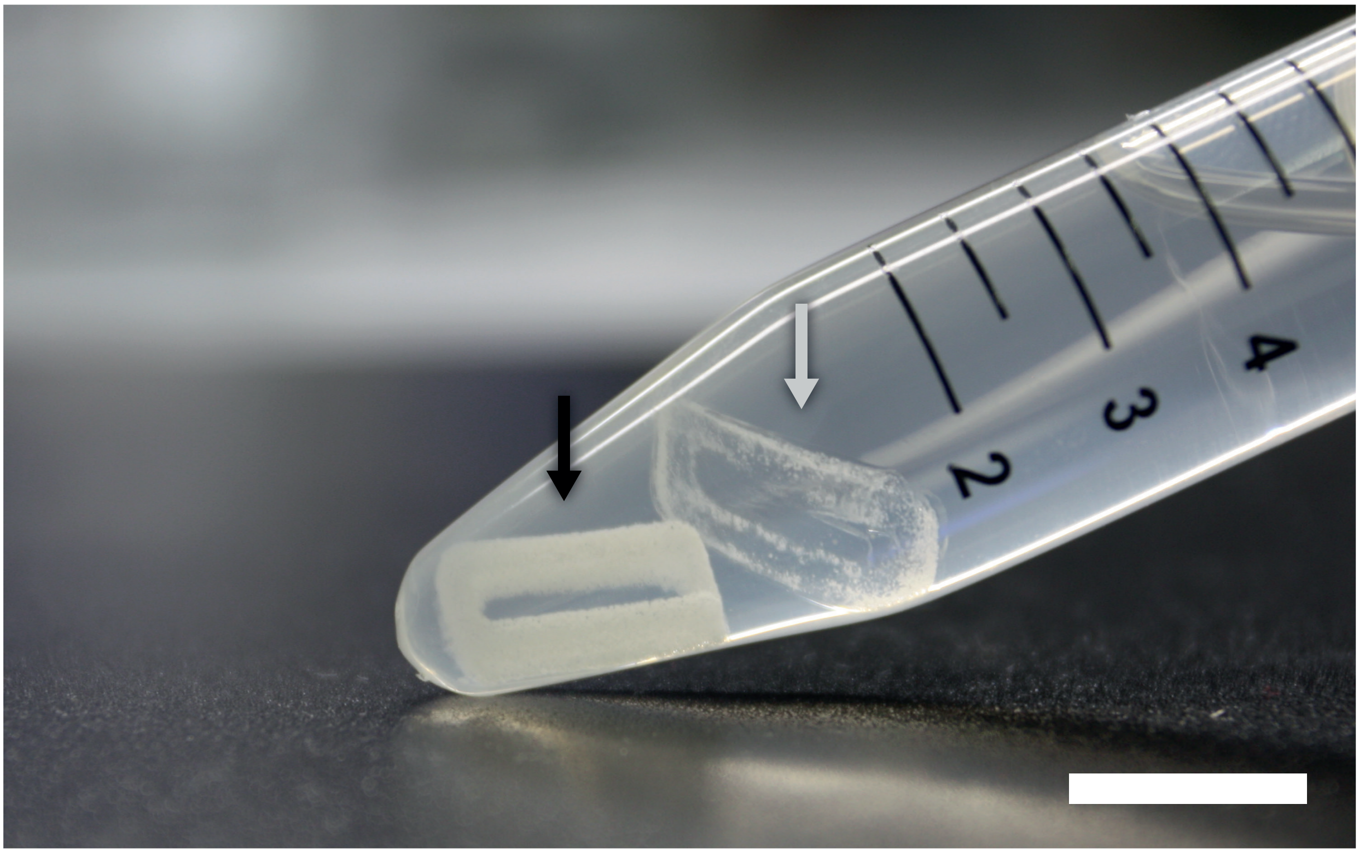

### Supplemental S7

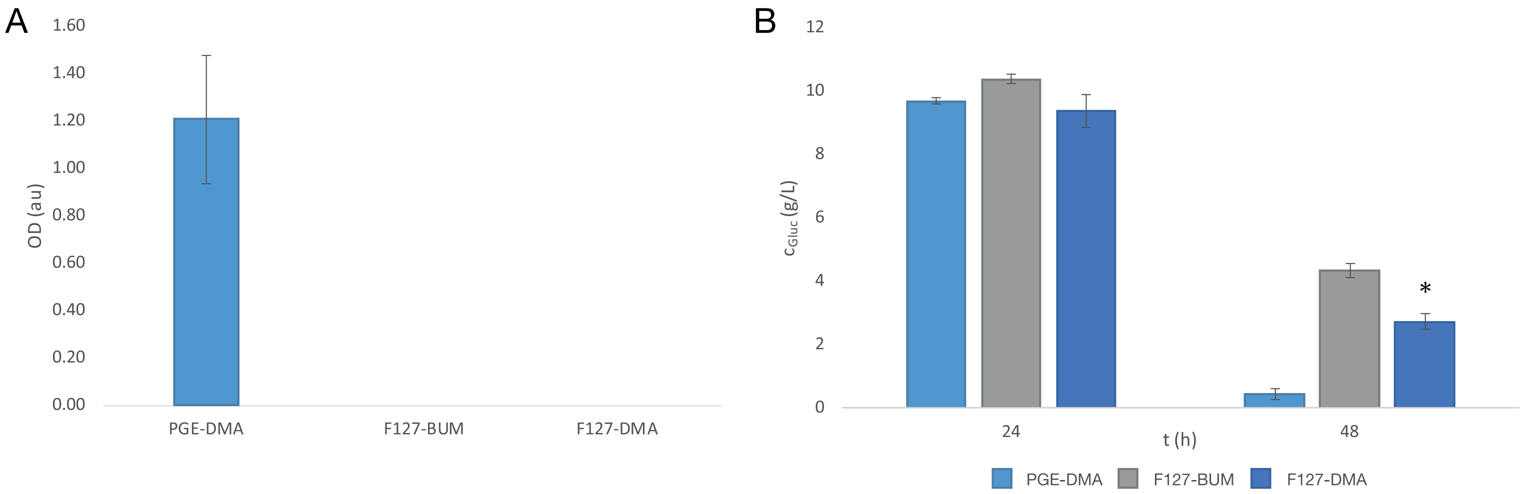

### Supplemental S8

Figure S8

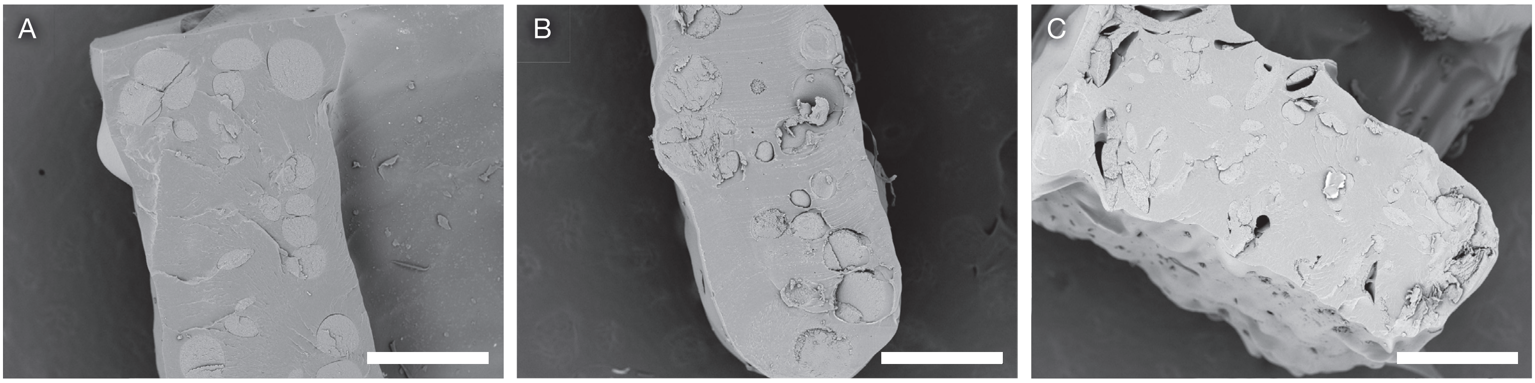

### Supplemental S9

Figure S9

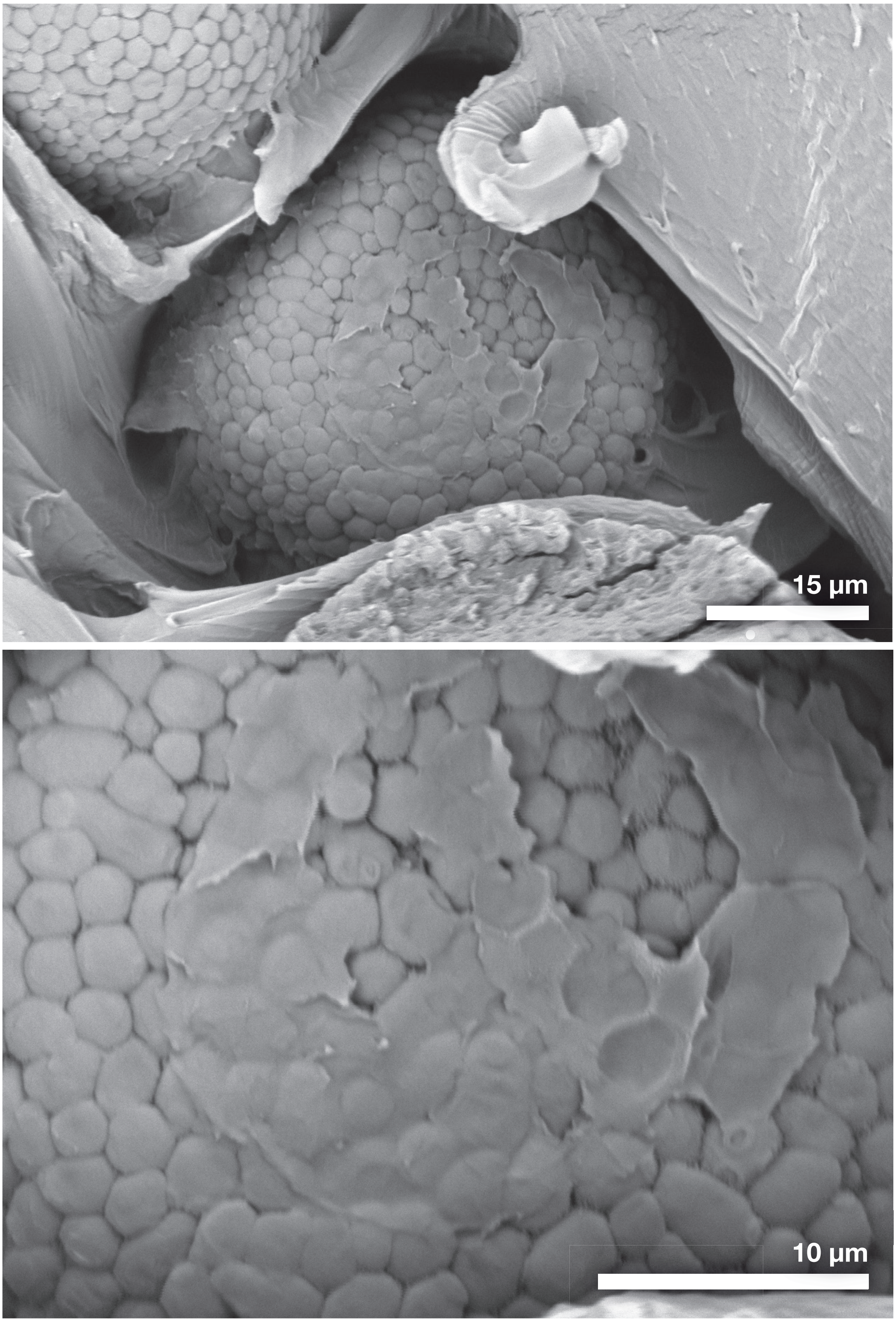

### Supplemental S10

Figure S10

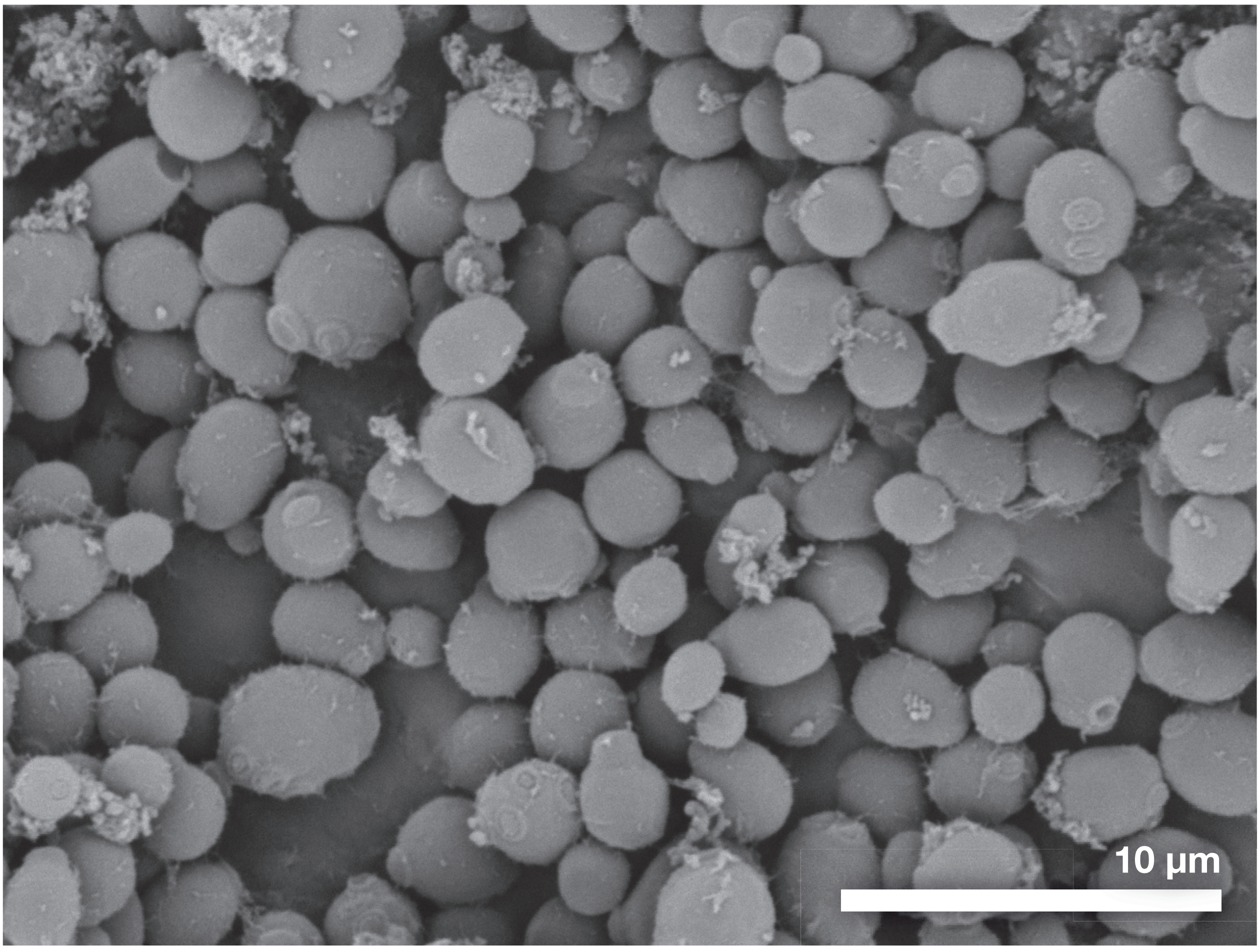
