## Supplemental S2 for "Physical confinement impacts cellular phenotype within living materials"

### Figure S2

A

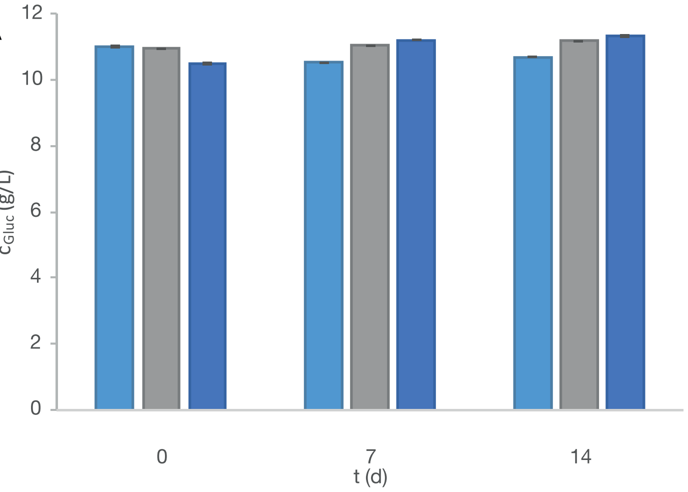


B

| Hydrogel | pH ($\log_{10}$ [$H^+$]) |
|---|---|
| PGE-DMA | 6.89 ± 0.03 |
| F127-BUM | 6.93 ± 0.01 |
| F127-DMA | 6.91 ± 0.02 |

C

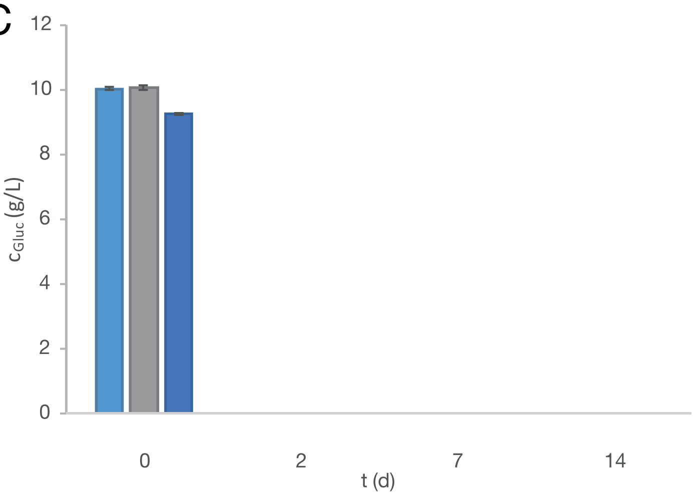


D

| Living material | pH ($\log_{10}$ [$H^+$]) |
|---|---|
| PGE-DMA | 5.90 ± 0.57 |
| F127-BUM | 6.30 ± 0.26 |
| F127-DMA | 6.13 ± 0.55 |

E

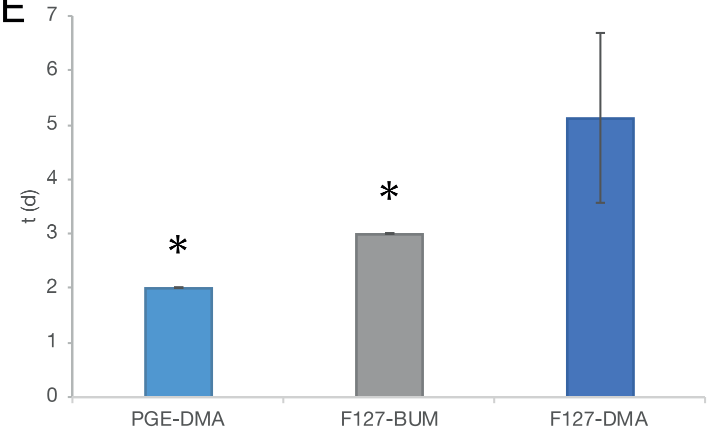


F

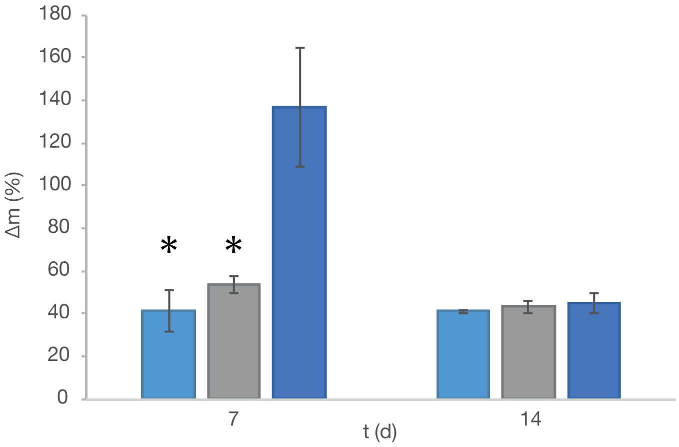


PGE-DMA F127-BUM F127-DMA
